# Decreased substrate stiffness leads to mitochondrial dysfunctions and Endothelial to Mesenchymal transition through Focal Adhesion Kinase activity in corneal endothelial cells

**DOI:** 10.1101/2025.11.08.687366

**Authors:** Sachin Anil Ghag, Viviane Souza de Campos, Subashree Murugan, Samuel Herberg, Rajalekshmy Shyam

**Author notes:** Corresponding author- 51 Newton Rd, Bowen Science Building 1-550, Iowa City, IA 52242.

## Abstract

**Purpose:** Fuchs’ Endothelial Corneal Dystrophy (FECD), a degenerative corneal disorder, is marked by the thickening of Descemet’s membrane and a progressive loss of corneal endothelial cells, ultimately leading to vision loss. A feature associated with the disease is the reduced stiffness of Descemet’s membrane. However, the effects of this change in Descemet’s membrane, on corneal endothelial cell health are not well understood. To explore this, we used *in vitro, in vivo,* and *ex vivo* studies to investigate how changes in substrate stiffness and the signaling pathways associated with these changes influence corneal endothelial functions.

**Methods:** For in-vitro studies, we cultured bovine corneal endothelial cells for 96 hours on stiff (32 kPa) and soft (8 kPa) substrate CytoSoft plates. By using Jess immunoassay and traditional western blotting, we evaluated changes in integrin signaling components, endothelial-to-mesenchymal transition, apoptosis, autophagy, and ubiquitin-proteasome pathway markers. Mitochondrial health and mitochondrial superoxide levels were assessed using commercial kits. We assessed the protein levels of the above-mentioned markers in the *Col8a2^Q455K/Q455K^* FECD mouse model. To evaluate whether FAK signaling contributes to the FECD pathogenesis, we injected the *Col8a2^Q455K/Q455K^* mice with an FAK inhibitor and assessed the corneal phenotypes.

**Results:** We observed increased levels of phosphorylated FAK, integrins α4 and α5 in bovine corneal endothelial cells cultured on soft substrate. We also found upregulated endothelial-to-mesenchymal transition (EndMT) markers, mitochondrial dysfunction, and apoptosis in cells grown on soft substrate. In the *Col8a2*^Q455K/Q455K^ mouse model of FECD, there was increased pFAK Y397 levels coincident with the onset of phenotypes. Intraperitoneal injections of a pFAK inhibitor improved antioxidant protein expression and decreased EndMT; however, it did not improve FECD-associated disease progression.

**Conclusion:** In this study, we explored how changes in the physical characteristics of the Descemet’s membrane impact corneal endothelial cell health. While we discovered activation of Focal adhesion kinase as a result of stiffness changes, its inhibition alone was insufficient to improve cell health in an FECD mouse model.

## Introduction

The corneal endothelium, a layer of non-proliferative cells, plays a vital role in preserving corneal transparency.^1^ Fuchs endothelial corneal dystrophy (FECD) is characterized by the degeneration of corneal endothelial cells and changes in Descemet’s membrane. Fuchs endothelial corneal dystrophy (FECD) is characterized by the degeneration of corneal endothelial cells and changes in Descemet’s membrane. There is no cure for this condition. Even though this disease affects one in twenty five adults over the age of 40 in the United States, current treatment is limited to corneal transplantation.^2,3^ Descemet’s membrane is an acellular layer composed of various extracellular matrix (ECM) proteins that provide structural support to the corneal endothelial cells. In turn, corneal endothelial cells secrete ECM components that are essential for both the formation and maintenance of this membrane.^4–6^ The ECM plays a key role in many cell types in regulating the biomechanical environment, strongly influencing how cells adhere, migrate, and function. Cells are therefore highly sensitive to changes in ECM composition or structure.^7–10^ Proulx *et. al.* recently discovered changes in corneal endothelial cell behavior because of the alterations in ECM components.^11^ As FECD progresses, the Descemet’s membrane stiffness reduces, largely due to abnormalities in its ECM composition.^4,12^ Atomic force microscopy studies report normal human Descemet’s membrane stiffness ranging from 20 to 80 kPa, with an average of 50 ± 17.8 kPa. In contrast, FECD-affected corneas show a markedly reduced stiffness, ranging from 2 to 15 kPa, with an average of 7.5 ± 4.2 kPa.^13^ Studies on animal models align with these findings. In wild-type mice, Descemet’s membrane stiffness averages 10.05 ± 7.34 kPa, whereas *Col8a2^Q455K/Q455K^*mutant mice- an early-onset FECD model- exhibit significantly lower stiffness at 4.37 ± 3.92 kPa.^2^ The impact of the stiffness changes to the Descemet’s membrane on corneal endothelial cells remains largely unexplored.

To fill this knowledge gap, we used *in vitro, in vivo,* and *ex vivo* studies to investigate how alterations in substrate stiffness affect corneal endothelial health.

## Materials and Methods

### Primary bovine corneal endothelial cell culture

Corneal endothelial cells were isolated from freshly obtained bovine eyes from local abattoirs. The cells were plated in T25 tissue culture flasks (Corning, cat#430168) with Dulbecco’s Modified Eagle Medium (Gibco, cat#11995065) supplemented with 10% Fetal Bovine Serum (Gibco, cat#10082-147) and 1x antibiotic-antimycotic (Gibco, cat#15240062). The cells were maintained at 37^0^C under 5% CO_2_ and then sub-cultured until passage 2 for experiments.

CytoSoft T25 flasks with stiffnesses of 32 kPa (Advanced Biomatrix, cat#5321-10EA) and 8 kPa (Advanced Biomatrix, cat#5319-10EA) were coated with 1 μg/ml fibronectin (Advanced Biomatrix, cat#5050) and 100 μg/ml Type-I Collagen (Advanced Biomatrix, cat#5005), diluted in 1X PBS, for 2 hours at room temperature according to the manufacturer’s instructions. The flasks were washed with the 1X PBS, and bovine corneal endothelial cells of passage 2 were seeded at a density of 2.1x10^6^ cells/cm² in the growth medium- DMEM + 10% Fetal Bovine Serum + 1% antibiotic/antimycotic as described elsewhere.^14,15^ The cells were cultured for 48 or 96 hours. After trypsinization, bovine corneal endothelial cells were washed twice with PBS and lysed using 1x RIPA buffer (Millipore Sigma, cat#20-188) containing protease and phosphatase inhibitors (Cell Signaling Technology, cat#5872s), 25 μM MG-132 (Cell Signaling Technology, cat#2194), and 50μM PR-619 (LifeSensors, cat# SI9619).^16^ The lysate was then centrifuged for 15 minutes at 13,000xg at 4 ^0^C. The supernatant was collected and used for either traditional Western blotting or capillary-based Jess immunoassay.

### Capillary-based Jess immunoassay

The Protein Simple Jess system, equipped with a 12-230 kDa separation module (SM-W002) was utilized according to the manufacturer’s instructions. The primary antibodies were used with 1:10 dilution for the Jess immunoassay. The data analysis was conducted using Compass for SW software, which provided electropherograms and virtual blot images. Normalization was carried out by using the total protein.

### Traditional Western blotting

25 µg of protein were then loaded onto either 12% or 15% SDS-PAGE gels for separation. After electrophoresis, the protein bands were transferred onto 0.22 µm nitrocellulose membranes (Li-COR, cat#926-31092). The membranes were subsequently incubated in blocking buffer (cat#927-60001) for 30 minutes, followed by overnight storage at 4^0^C with primary antibodies-LC3, SQSTM1/p62 (1:1000 dilution) in antibody diluent (Li-COR, cat#65001). The next day, membranes were washed in Tris Base buffer containing 0.1% tween 20 and incubated in secondary antibodies- Rabbit (cat#926-68071) or mouse (cat#926-32210) for 1 hour at room temperature. Visualization of the blots was achieved using the Odyssey XF imager (Li-COR, cat#2802-01). Protein expressions were normalized to total protein stain (Li-COR, cat#926-10016) levels.

### Mice

C57BL/6 FECD mice carrying the *Col8a2*^Q455K/Q455K^ knock-in mutation were purchased from the Jackson Laboratory (strain # 029749, Bar Harbor, ME).^17^ We used 7- and 10-week-old *Col8a2*^Q455K/Q455K^ knock-in mice, along with age-matched wild-type controls, for all experiments. Genotyping was performed through an automated service (Transnetyx, Cordova, TN). All the experimental research in the current study has been approved by the Institutional Animal Care and Use Committee (IACUC) at Indiana University and adhered to the Association for Research in Vision and Ophthalmology (ARVO) statement for the use of animals in Ophthalmic and Vision research.

### Intraperitoneal Injections, Optical Coherence Tomography (OCT), and Heidelberg Retinal Tomography3 (HRT)

7-week-old FECD mice carrying the Col8a2^Q455K/Q455K^ knock-in mutation were injected intraperitoneally with p-FAK inhibitor- Ifebemtinib (BI853520, MedChemExpress, cat#HY-122844). 5.88 mg/kg (10 μM/kg) of Ifebemtinib dissolved in DMSO was injected once a day for 2 weeks.^18,19^ Mice injected with 0.1% DMSO were used as controls. Anterior segment- Optical Coherence Tomography (AS-OCT) (iVue100 Optovue, Inc., Fremont, CA, USA) and Heidelberg Retinal Tomography 3-Rostock Cornea Module (HRT3-RCM) (Heidelberg Engineering Inc., Franklin, MA, USA) were performed to measure the corneal thickness and assess the corneal endothelial cells post-treatment.

### Mice Corneal Endothelial Collection

Eyes enucleated from mice were stored in ice-cold 1× PBS. The corneal endothelium, along with Descemet’s membrane, was carefully dissected and peeled away from the cornea. For each sample, six corneal endothelia from three mice were pooled to prepare a protein lysate. Tissues were lysed in 1× RIPA buffer (Millipore Sigma, Cat# 20-188) supplemented with protease and phosphatase inhibitors (Cell Signaling Technology, Cat# 5872S), 25 μM MG-132 (Cell Signaling Technology, Cat# 2194), and 50 μM PR-619 (LifeSensors, Cat# SI9619). Lysates were centrifuged at 13,000 × g for 15 minutes at 4°C, and the resulting supernatant was collected for analysis using a capillary-based Jess immunoassay.

### Chymotrypsin activity assay

The activity of chymotrypsin-like protease was quantified using a commercially available Amplite Fluorometric Proteasome 20S Activity Assay Kit (AAT Bioquest, cat#13456). This kit employs LLVY-R110 as a fluorogenic substrate, which emits green fluorescence upon cleavage by the proteasome. According to the manufacturer’s protocol, bovine corneal endothelial cells (0.02 x 10^6^ cells per well) were grown in 100 μL of culture media in 96-well CytoSoft plates coated with type-I collagen and fibronectin for 48 and 96 hours. Following the incubation in the assay working solution for 1 hour at 37^0^C, fluorescence emissions were recorded using a microplate reader (Biotek-Synergy-HTX), using an excitation wavelength range of 480-500 nm, and emission wavelength range of 520-530 nm.

### JC-1 staining

Bovine corneal endothelial cells (0.02 x 10^6^ cells per well) were cultured on 96-well CytoSoft plates coated with type-I collagen and fibronectin for 48 and 96 hours. The cell culture medium was removed and the cells were washed with warm (37^0^C) 1X PBS, after which fresh warmed cell culture medium was added. Subsequently, JC-1 dye (Invitrogen, cat#T3168) was added to achieve a final concentration of 2 μM, and the cells were incubated at 37^0^C in a 5% CO_2_ environment for 20 minutes. A final concentration of 50 μM CCCP (Thermo Fisher, cat#M34152) was introduced for the positive control, and the cells were incubated at 37^0^C for 5 minutes. After incubation, the culture medium from all samples was discarded, and cells were washed once with warm 1X PBS. 100 μl 1X PBS for each well of the 96-well plate was added to the samples. The fluorescence was analyzed immediately in a microplate reader (Biotek-Synergy-HTX) using an excitation wavelength range of 480-500 nm, and an emission wavelength range of 580-590 nm.

### Mitosox assay

To assess Mitochondrial Superoxide, the commercially available MitoSOX™ Kit (Thermo Fisher, cat#M36008) was used, following the manufacturer’s instructions. bovine corneal endothelial cells were seeded at a density of 0.02 x 10^6^ cells per well on 96-well CytoSoft plates coated with type-I collagen and fibronectin for 48 and 96 hours. Cells were stained with 1 μM MitoSOX Red for 30 minutes at 37 ^0^C. The fluorescence was assessed using a microplate reader (Biotek-Synergy-HTX) at an excitation wavelength range of 480-500 nm, and an emission wavelength range of 580-590 nm.

### Statistical analysis

Data was analyzed using GraphPad Prism 10.0.1. All data are shown with mean ± standard deviation. The unpaired Student’s t-test and 2-way ANOVA with Tukey’s multiple comparisons were used for statistical analysis with GraphPad Prism 8.0 software. A p-value of 0.05 or lower was considered statistically significant.

## Results

### Change in substrate stiffness upregulates focal adhesion kinase and integrins

Since the decrease in Descemet’s membrane stiffness is associated with FECD,^2,13^ we assessed whether growing normal primary corneal endothelial cells on soft substrates will change the expression of key integrins and activate the integrin signaling transducer, Focal adhesion kinase (FAK).^20^ Bovine corneal endothelial cells are a well-established alternate for human cells.^14,21^ When cultured in soft (8 kPa) substrate, primary bovine corneal endothelial cells exhibited a notable increase in the phosphorylation of FAK Tyr 397 site (p=0.002, n=3) at 96 hours compared to cells cultured on stiff (32 kPa) substrates (**Fig. 1A and B**). Additionally, integrins α4 and α5 showed significant upregulation on the soft substrate after 96 hours (p = 0.008 for both), compared to the stiff substrate (**Fig. 1C and D**), indicating activation of the FAK-integrin pathway activity. In contrast, we observed decreased expression of integrin β4 (p=0.01) and no changes in the levels of integrin β3 (**Fig. 1C and D**).

**Figure 1.**
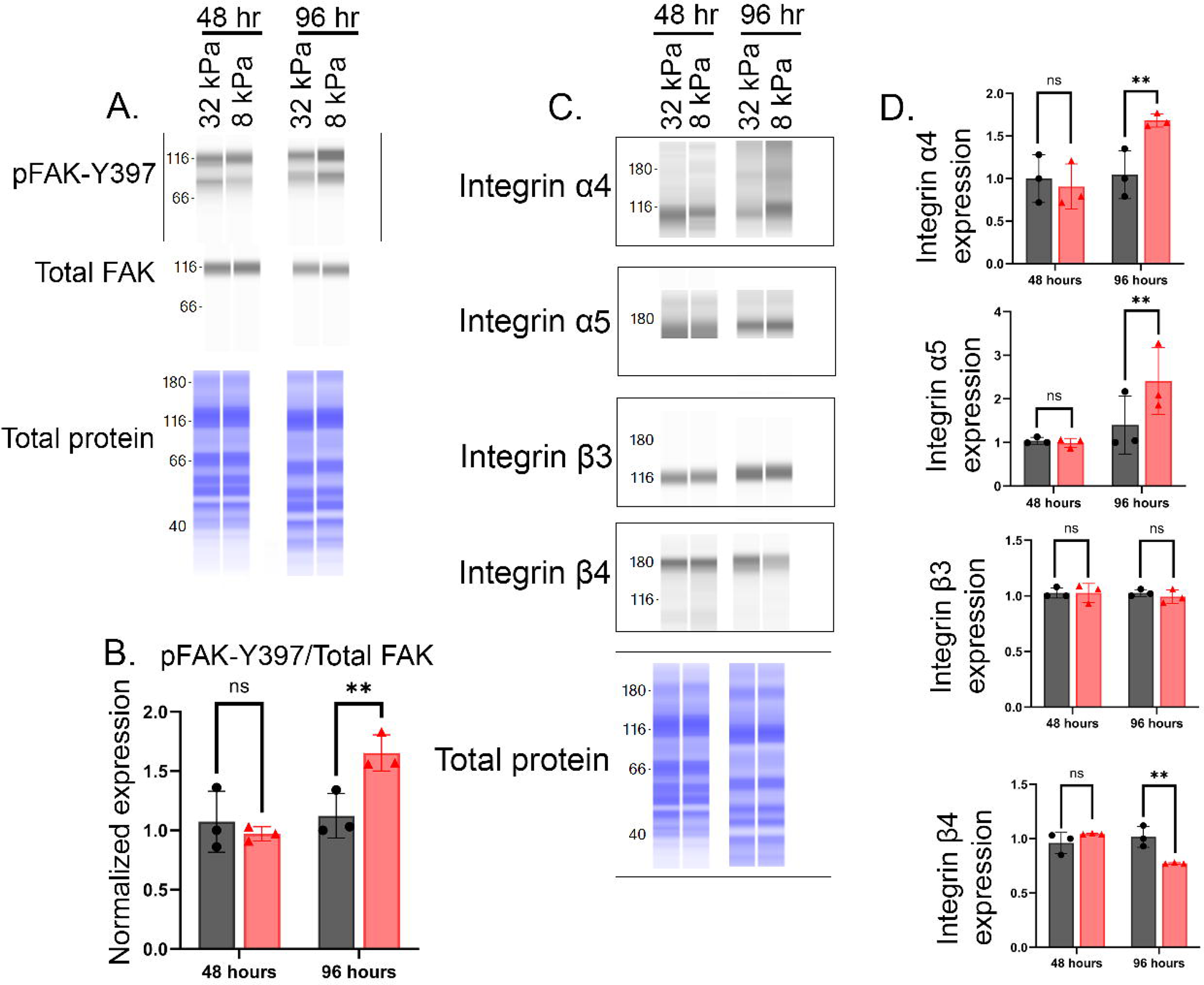
Substrate stiffness regulates FAK activation and integrins expression in cultured cells. **(A)** Representative Jess immunoassay blots for phosphorylation at tyrosine 397-Focal Adhesion Kinase (p-FAK-Y397) and total FAK (t-FAK) in bovine corneal endothelial cells cultured on stiff (32 kPa) or soft (8 kPa) substrates mimicking healthy and FECD-like conditions, respectively, for 48 and 96 hours. **(B)** Quantification of blots for panel A normalized by total protein; Mean ± standard deviation; n=3; unpaired Student’s t-test; ns (not significant), **p < 0.01. **(C)** Representative Jess immunoassay blots for integrins α4,α5,α6,β3, and β4. **(D)** Quantification of blots for panel-C normalized by total protein; Mean ± standard deviation; n=3; unpaired Student’s t-test; ns (not significant), **p < 0.01.

### Decreased substrate stiffness results in endothelial-to-mesenchymal transition and cell death in bovine corneal endothelial cells

Increased expression of key endothelial-to-mesenchymal transition (EndMT) genes, ZEB1 and Snail, is associated with FECD.^22^ We investigated whether soft substrate was sufficient to induce EndMT in primary bovine corneal endothelial cells. Bovine corneal endothelial cells cultured on 8 kPa plates showed increased expression of Snail (p=0.02, n=3), ZEB-1(p=0.009, n=3)(**Fig. 2A and C**). Apoptotic cell death is present in FECD corneal endothelial cells.^23,24^ Therefore, we evaluated whether a soft substrate could cause apoptosis in healthy primary cells. We found significant upregulation in the ratio of cleaved caspase 9 and total caspase 9 (p=0.035, n=3) at 96 hours (**Fig. 2B and D**).

**Figure 2.**
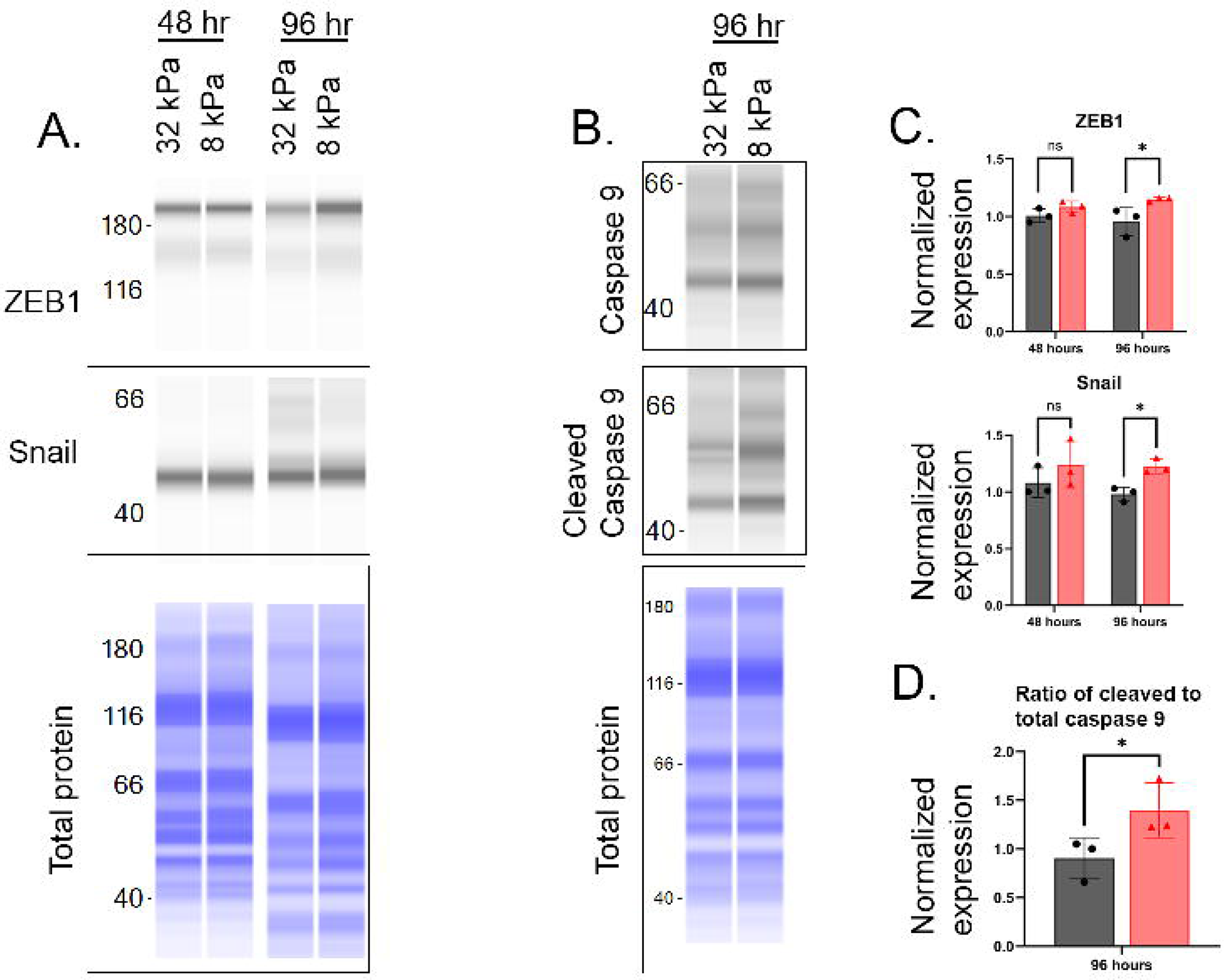
Matrix stiffness promotes endothelial to mesenchymal transition-associated proteins and apoptotic proteins expression. **(A)** Representative Jess immunoassay blots for endothelial to mesenchymal transition markers- ZEB1 (Zinc finger E-box binding homeobox 1) and Snail (Zinc finger protein SNAI1) in bovine corneal endothelial cells cultured on stiff (32 kPa) or soft (8 kPa) substrates for 48 and 96 hours. **(B)** Representative Jess immunoblots for apoptosis markers- cleaved caspase 9 and total caspase 9 in bovine corneal endothelial cells cultured in different stiffness (32 kPa and 8 kPa) for 96 hours. **(C)** Quantification of blots for panel A normalized by total protein; Mean ± standard deviation; n=3; unpaired Student’s t-test; ns (not significant), *p < 0.05 **(D)** Corresponding quantification of the panel B normalized by total protein; Mean ± standard deviation; n=3; unpaired Student’s t-test; ns (not significant), *p<0.05.

### Mitochondrial dysfunction in bovine corneal endothelial cells cultured in 8 kPa substrate

Increased mitochondrial dysfunctions are strongly associated with FECD pathogenesis.^25–27^ We analyzed whether substrate stiffness can negatively impact mitochondrial health. The protein expression analysis (**Fig. 3A and B**) did not show any change in the expression of mitochondrial antioxidant, SOD2, and overall antioxidant, Catalase. Interestingly, SLC4A11- a mitochondrial proton uncoupler crucial for corneal endothelial functions^25^ - was increased in cells cultured on the 8 kPa substrate (**Fig. 3A and B**).

**Figure 3.**
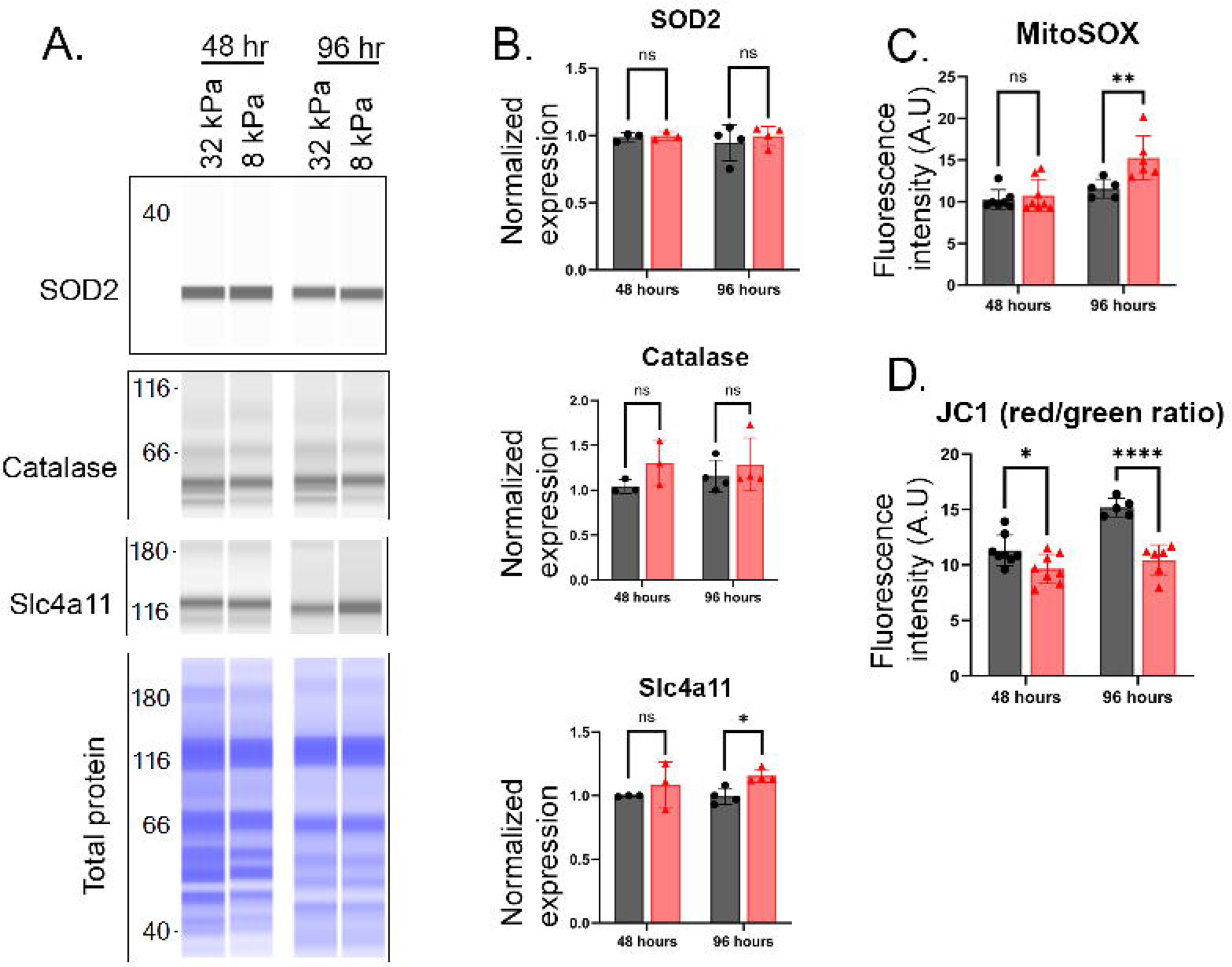
Substrate stiffness alters mitochondrial homeostasis and oxidative stress response. **(A)** Representative Jess immunoblots for Superoxide dismutase 2, mitochondrial (SOD2), catalase, and solute carrier family 4 member 11 (SLC4A11) in bovine corneal endothelial cells cultured on stiff (32 kPa) or soft (8 kPa) substrates for 48 and 96 hours. **(B)** Corresponding quantification of the blots normalized by total protein; Mean ± standard deviation; n=3-4; unpaired Student’s t-test; ns (not significant), *p<0.05. **(C)** Mean intensity of MitoSOX fluorescence quantified in 32 kPa versus 8 kPa to measure mitochondrial superoxide levels. Mean ± standard deviation; n=5-8; unpaired Student’s t-test; ns (not significant), **p<0.01 **(D)** Mean fluorescence intensities of red and green emission ratios quantified after JC-1 staining. Mean ± standard deviation; n=5-8; unpaired Student’s t-test; ns (not significant), *p<0.05, ****p < 0.0001.

We evaluated mitochondrial integrity and reactive oxygen species production in bovine corneal endothelial cells cultured on 32 kPa and 8 kPa substrates using commercial kits. We observed increased mitochondrial superoxide production in cells grown on the 8 kPa substrate when compared to those grown on 32 kPa (p=0.003, n=5) (**Fig. 3C**). Shift in JC-1 emission from red to green indicates depolarization of mitochondria. Bovine corneal endothelial cells grown on soft substrate showed increased mitochondrial depolarization (**Fig. 3D**). Accordingly, the data showed a decreased red (∼590 nm) to green (∼529 nm) fluorescence intensity ratio in 8 kPa plates compared to 32 kPa (p=0.016, n=8 at 48 hours and p=>0.0001, n=6 at 96 hours). These findings suggest that substrate stiffness influences reactive oxygen species generation without markedly affecting mitochondrial antioxidant levels.

### Proteasome activity increased in bovine corneal endothelial cells cultured in 8kPa plates

Previous studies demonstrated enhanced autophagy in FECD,^28,29^ and our recent studies show that decreased proteasome activities can lead to FECD-associated features in healthy mice.^16^ Therefore, we assessed whether ECM stiffness can affect these protein degradation pathways in bovine corneal endothelial cells. We first evaluated the expression of the autophagy marker proteins LC3 and p62. LC3 exists in cytosolic (LC3-I) and membrane-bound (LC3-II) forms; the ratio of the latter increases with the formation of autophagosomes. p62 acts as a cargo receptor for autophagy, binding to ubiquitinated proteins and LC3. We did not observe any changes in the expression of these proteins with alterations in substrate stiffness (**Fig. 4A and B**). However, we noted a significant increase in the chymotrypsin activities of proteasomes (p=0.0001 at 48 hours; p=0.03, n=3 at 96 hours) in the cells grown on 8 kPa substrates compared to 32 kPa (**Fig. 4C**). Among the ubiquitin proteasome pathway proteins, we observed increased levels of 26S proteasome subunit, PSMD11. However, we did not see any changes in K48 ubiquitylation, a proteasome-specific ubiquitin linkage and 11S regulator, PA28α expression in soft substrates (**Fig. 4D and E**). Together, these results suggest that soft ECM conditions enhance proteasomal activity but do not increase the expression of proteasome associated proteins or autophagy in bovine corneal endothelial cells.

**Figure 4.**
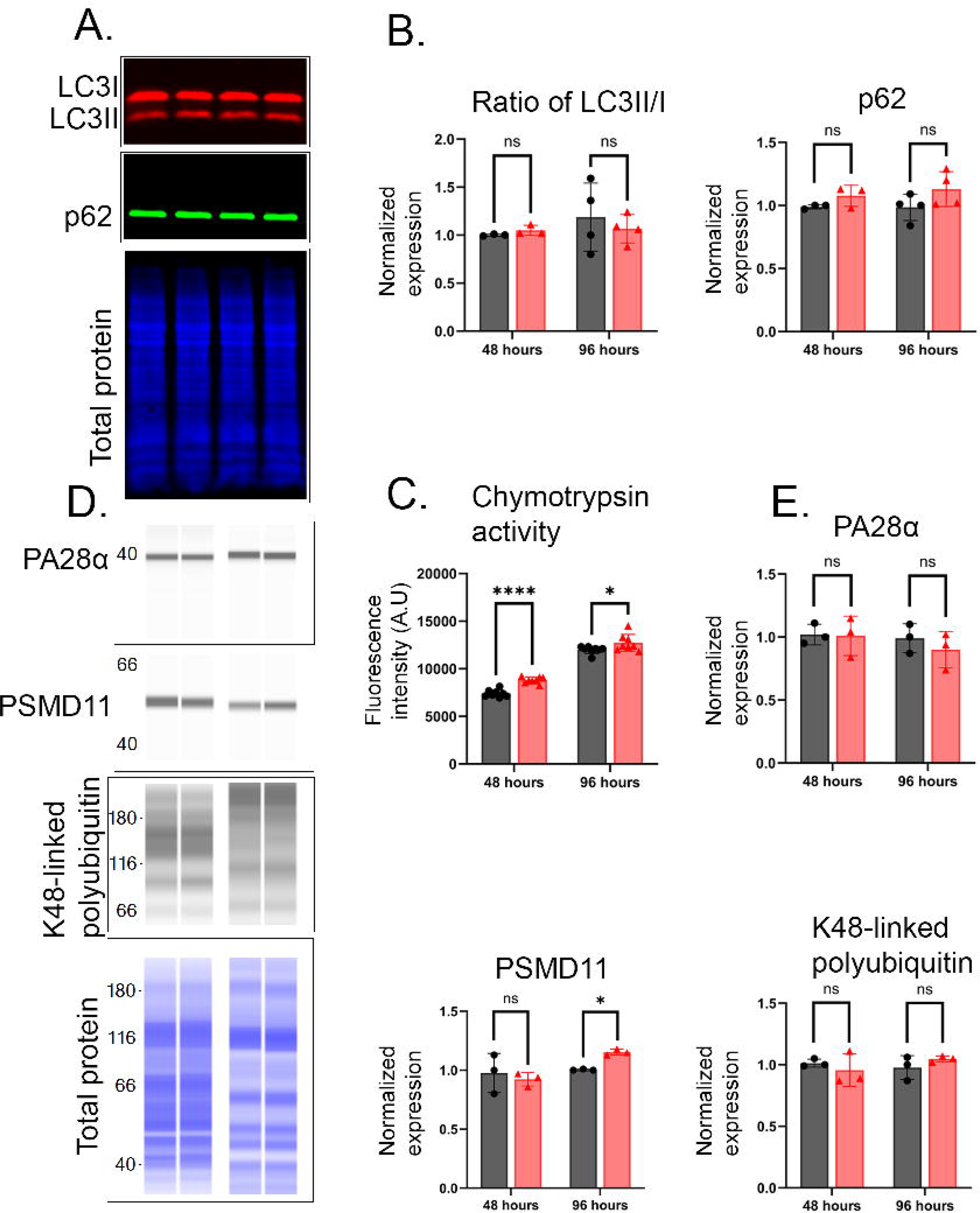
Substrate stiffness increased proteasomal activity and PSMD11 expression, but not autophagy-associated proteins or proteasome subunit levels. **(A)** Representative western blots for autophagy markers- Microtubule-associated protein light-chain 3 (LC3) and Sequestosome 1 (p62/SQSTM1) in bovine corneal endothelial cells cultured on stiff (32 kPa) or soft (8 kPa) substrates for 48 and 96 hours. **(B)** Corresponding quantification of the blots normalized by total protein; Mean ± standard deviation; n=3-4; unpaired Student’s t-test; ns (not significant). **(C)** Quantification of chymotrypsin activity, Mean ± standard deviation; n=8-10; unpaired Student’s t-test; ns (not significant), *p<0.05, ****p < 0.0001. **(D)** Representative Jess immunoblots for ubiquitin proteasomal pathway markers- proteasome 11S regulator subunit REG alpha (PA28α), proteasome 26S subunit non-ATPase 11 (PSMD11), and K48-linked poly-ubiquitin. **(E)** Corresponding quantification of the blots normalized by total protein; Mean ± standard deviation; n=3; unpaired Student’s t-test; ns (not significant), *p<0.05.

### Elevated focal adhesion kinase and integrins in a FECD mouse model

*Col8a2^Q455K/Q455K^* animals show FECD-associated features such as corneal endothelial cell loss at 5 weeks of age and prominent guttae at 10 weeks of age.^17^ These animals also show a significant decrease in Descemet’s membrane stiffness at 5 weeks.^2^ In the *Col8a2^Q455K/Q455K^* animals, at the onset of disease-associated phenotypes (10-week-old), we observed increase in the expression of p-FAK-Y397 (p=0.008, n=3) and integrin α4 (p=0.114, n=3) expression (**Fig. 5A-D**), which are consistent with *in vitro* findings. Similar to the *in vitro* study, we found a significant upregulation of EndMT markers- ZEB1 (p=0.022, n=3) and Snail (p=0.012, n=3) in mice (**Fig. 5E and F**).

**Figure 5.**
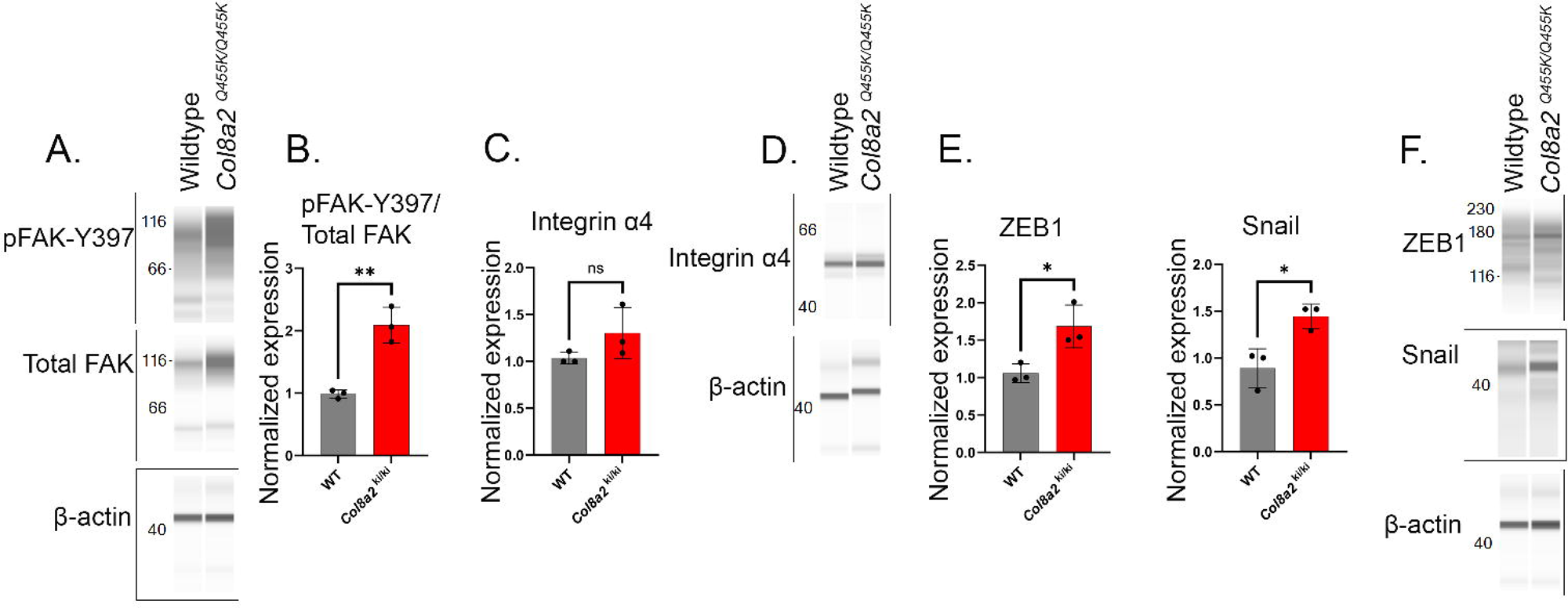
Fuchs’ Endothelial Corneal Dystrophy (FECD) mouse model exhibits corneal endothelial alterations that are associated with changes in substrate stiffness. **(A)** Representative blots from JESS immunoassay of 10-week-old FECD mice (*Col8a2^Q455K/Q455K^*) versus age-matched wildtype mice (WT) for p-FAK-Y397 and total FAK. **(B)** Quantification of the Jess immunoblots in panel A normalized to β-Actin. Mean ± standard deviation; n=3; unpaired Student’s t-test; ns (not significant), *p < 0.05, **p < 0.01. **(C and D)** Representative blots from JESS immunoassay of FECD versus WT mice for integrin α4. Quantification of the Jess immunoblots in panel C normalized to β-Actin. Mean ± standard deviation; n=3; unpaired Student’s t-test; ns (not significant). **(E and F)** Representative blots from JESS immunoassay of FECD versus WT mice for endothelial to mesenchymal transition markers- ZEB1 (Zinc finger E-box binding homeobox 1), Snail (Zinc finger protein SNAIL). Quantification of the Jess immunoblots in panel G normalized to β-Actin. Mean ± standard deviation; n=3; unpaired Student’s t-test; ns (not significant), *p < 0.05.

### Inhibition of phosphorylated FAK increases the expression of apoptotic markers, as well as antioxidant enzymes and SLC4A11, in the corneal endothelium of FECD mice

To understand the association between upregulation of FAK phosphorylation and FECD progression, we injected 7-week-old Col8a2^Q455K/Q455K^ knock-in FECD mice with a highly potent, selective ATP-competitive FAK inhibitor- Ifebemtinib.^18^ It inhibits FAK autophosphorylation at tyrosine 397, which also inhibits downstream signaling pathways such as Src and PI3K/AKT.^18^ Two weeks post-treatment, mice injected with Ifebemtinib showed no significant changes in body weight and intra-ocular pressure (n=18) (**Fig. 6A and B**). OCT measurements revealed no change in the corneal thickness (p=0.295, n=18) (**Fig. 6C and D**). However, HRT imaging revealed a significant decrease in corneal endothelial cell density (p=0.0009, n=18). (**Fig. 6F**).

**Figure 6.**
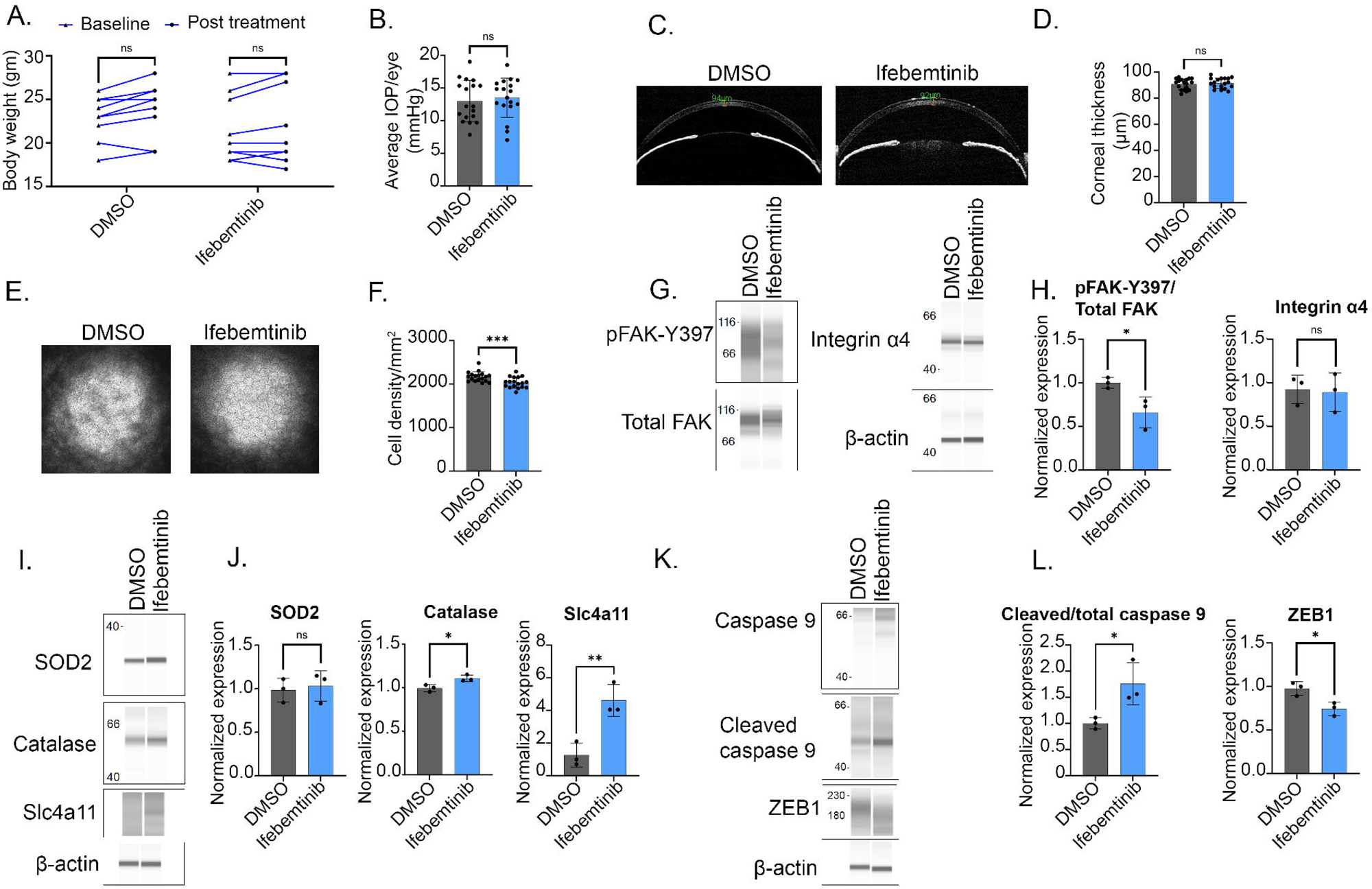
Pharmacological inhibition of pFAK downregulates endothelial to mesenchymal transition, upregulates antioxidants and SLC4A11 levels, but also upregulates apoptotic markers in Fuchs Endothelial Corneal Dystrophy (FECD) mouse. **(A)** Longitudinal measure of the body weight in control and Ifebemtinib-injected mice at baseline, 2-weeks post-treatment. Mean ± standard deviation; n=9; 2-way ANOVA with Tukey’s multiple comparisons; ns (not significant). **(B)** Intra-ocular pressure (IOP) measurement in control and Ifebemtinib-injected mice at 2-weeks post-treatment. Mean ± standard deviation; n=18 eyes; unpaired Student’s t-test; ns (not significant). **(C)** Representative OCT images of control and Ifebemtinib-injected mice post-treatment. **(D)** Quantification of the corneal thickness in control and Ifebemtinib-injected mice post-treatment. Mean ± standard deviation; n=18 eyes; unpaired Student’s t-test; ns (not significant). **(E)** Representative HRT3 images of control, Ifebemtinib-injected mice at 2 weeks post-treatment. **(F)** Quantification of the endothelial cell density/mm^2^ in control and Ifebemtinib-injected mice post-treatment. Mean ± standard deviation; n=18 eyes; unpaired student’s t-test; ns (not significant), ***p < 0.001. **(G)** Representative blots from JESS immunoassay of control versus Ifebemtinib-injected mice for p-FAK-Y397, total FAK and integrin α4. **(H)** Quantification of the Jess immunoblots in panel G normalized to β-Actin. Mean ± standard deviation; n=3; unpaired Student’s t-test; ns (not significant), *p < 0.05, **p < 0.01. **(I)** Representative blots from JESS immunoassay of control versus Ifebemtinib-injected mice for Superoxide dismutase 2, mitochondrial (SOD2), catalase, and solute carrier family 4 member 11 (SLC4A11). **(J)** Representative blots from JESS immunoassay of control versus Ifebemtinib-injected mice for integrin α4. **(J)** Quantification of the Jess immunoblots in panel I normalized to β-Actin. Mean ± standard deviation; n=3; unpaired Student’s t-test; ns (not significant). **(K)** Representative blots from JESS immunoassay of control versus Ifebemtinib-injected mice for endothelial to mesenchymal transition marker- ZEB1 (Zinc finger E-box binding homeobox 1), apoptosis markers- cleaved caspase 9 and total caspase 9. **(L)** Quantification of the Jess immunoblots in panel K normalized to β-Actin. Mean ± standard deviation; n=3; unpaired Student’s t-test; ns (not significant), *p < 0.05.

**Figure 7.**
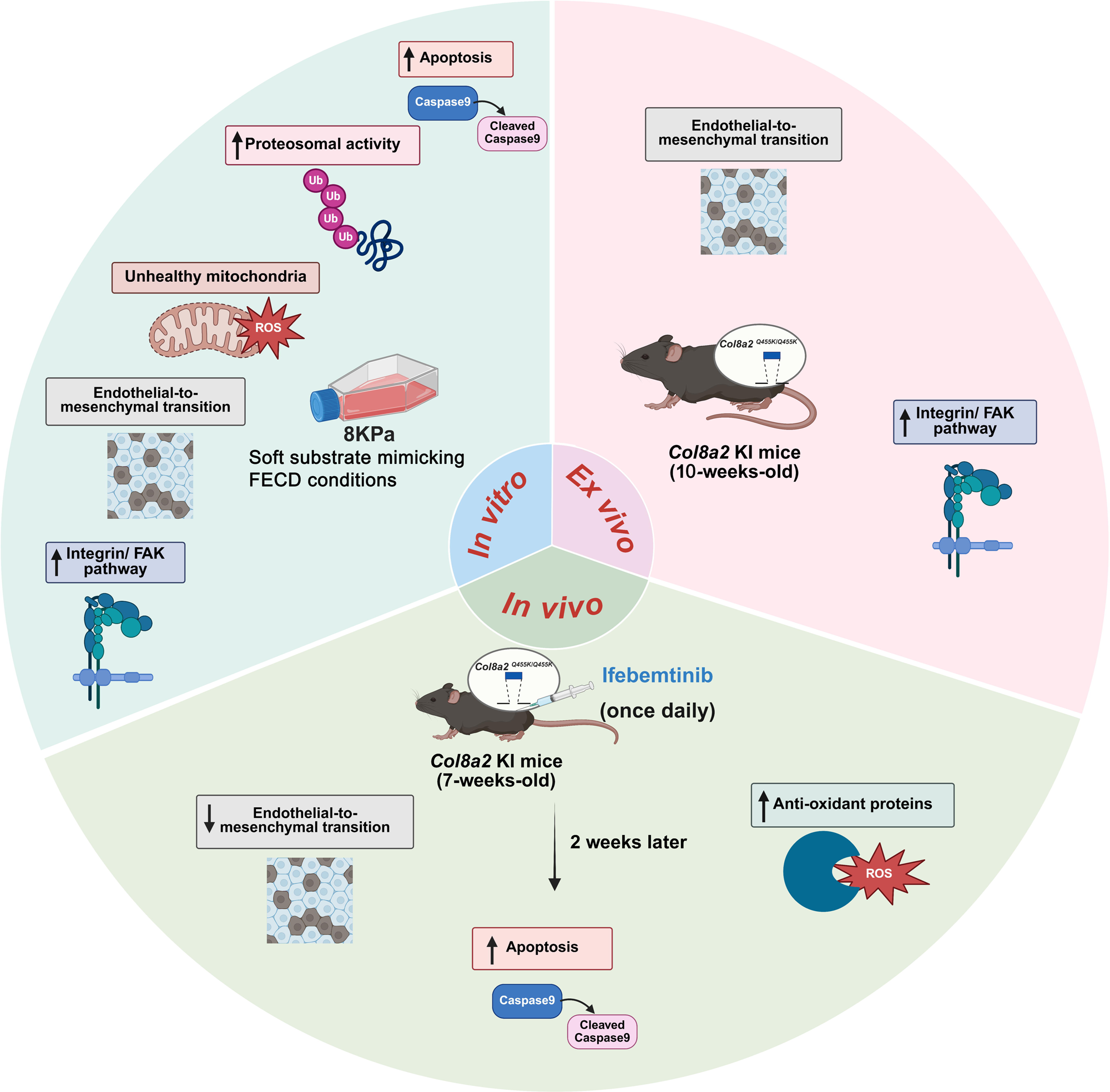
Graphical representation of the consequences of soft substrate stiffness on corneal endothelial health.

Jess immunoassay showed a significant downregulation in the phosphorylation of FAK (p=0.033, n=3) and integrin α4 (p=0.421, n=3) (**Fig. 6G and H**) in Ifebemtinib-injected mice corneal endothelial peelings confirming the efficacy of the treatment. Next, we assessed the effect of FAK inactivity on mitochondrial health. We observed a significant upregulation in SLC4A11 expression (p=0.005, n=3) and antioxidant protein, catalase (p=0.012, n=3), while no significant change in SOD2 (p=0.367, n=3) levels (**Fig. 6I and J**).

Finally, consistent with the changes in cell density with Ifebemtinib treatment, we also observed a significant increase in apoptotic cell death, i.e., an increase in the ratio of cleaved caspase 9 and total caspase 9 (p=0.030, n=3) in Ifebemtinib-injected mice compared to vehicle control (**Fig. 6K and L**). Also, we found decrease in the EndMT marker, ZEB1 (p=0.047, n=3) in Ifebemtinib-injected mice when compared to vehicle control mice (**Fig. 6K and L**).

## Discussion

Decreased Descemet’s membrane stiffness is associated with FECD progression.^27^ In this study, we evaluated how this change can impact corneal endothelial functions. We found that growing cells on a soft substrate was sufficient to induce FAK phosphorylation. While expression of integrin pathway components in the corneal endothelium during FECD is highlighted in many studies,^30,31^ our work provides evidence that physical changes to the substrate are sufficient to induce increased integrin expression and FAK phosphorylation. Understanding how alterations in stiffness influence corneal endothelial cells provides valuable insights into the extrinsic mechanisms driving FECD. It could help guide the development of therapies aimed at restoring normal corneal function.

Corneal endothelial cells are highly metabolic. Therefore, mitochondrial activities are essential for the normal functions of these cells.^15,23^ Dysfunctional mitochondria in FECD are known to be a significant source of reactive oxygen species.^29,32^ ECM changes can influence mitochondrial functions,^33^ and we saw such effects in the corneal endothelial cells, too. Decrease in substrate stiffness resulted in increased mitochondrial oxidative stress and depolarization, coincident with the increase in FAK phosphorylation. Inhibiting FAK activity restored mitochondrial antioxidant levels, suggesting a role of this pathway in regulating mitochondrial functions in the corneal endothelium.

SLC4A11 is an important protein in the corneal endothelium, aiding in the maintenance of mitochondrial health and plasma membrane integrity.^25,34,35^ SLC4A11 expression is decreased in FECD corneal endothelia.^36–38^ By growing cells on substrate stiffness similar to that of FECD corneas, we expected decreased expression of SLC4A11. Our observations, contrary to our predictions, suggest that while SLC4A11 protein expression is highly sensitive to substrate changes, the alteration of substrate physical properties might not be the reason behind the decreased expression of SLC4A11 in FECD.

ECM accumulation and activation of EndMT are commonly seen in FECD cells.^22,39,40^ Cells typically sense changes in ECM characteristics, such as composition and stiffness, through integrin transmembrane receptors. Altered ECM-integrin signaling can change cellular behavior and promote EndMT.^20,41^ Elevated expression of key EndMT regulators such as ZEB1 and Snail has been linked to FECD progression.^22^ In our study, we observed that softer substrates led to increased expression of ZEB1 and Snail. We observed that substrate stiffness has a significant impact on the morphological characteristics of bovine corneal endothelial cells. These findings suggest that substrate stiffness selectively influences EndMT. The pathways regulating these transitions will be explored in future studies.

We assessed the effect of altered substrate on autophagy and the ubiquitin proteasome pathway, the major protein clearance pathways.^16,42^ In FECD tissues, increased expression of several autophagy-related genes has been reported.^43^ Accumulation of p62 often signals impaired autophagic flux and has been linked to multiple degenerative disorders, including FECD.^42,44^ Although the LC3-II/I ratio did not significantly differ, we also did not observe increased p62 levels at 96 hours, suggesting decreased substrate stiffness may not impact autophagy. However, without studying autophagy flux, it is not possible to conclude the lack of influence of substrate stiffness on this pathway. We previously discovered that decreased proteasome activities can lead to the development of FECD-associated symptoms.^16^ Contrary to our expectations, we observed elevated chymotrypsin activity and an increase in 26S proteasome subunit, PSMD11, in cells grown on 8 kPa substrates. We also saw no changes to two additional markers of proteasome activity in these conditions.

In a mouse model of FECD, we observed a significant increase in phosphorylated FAK (p-FAK-Y397) along with its upstream regulator, integrin α4, indicating activation of the integrin–FAK signaling. These changes suggest that alterations in the physical environment may drive FAK-dependent signaling in FECD endothelial cells. We also detected higher levels of the EndMT markers ZEB1 and Snail, further supporting the idea that aberrant activation of EndMT contributes to FECD pathology.

FAK is well recognized for its role in promoting cell survival and protecting against apoptosis. In adherent cells, this effect is largely mediated through integrin-dependent pathways.^45,46^ Also, low expression of FAK can evoke an apoptotic response.^47^ Interestingly, increased death in cells grown on 8 kPa substrate, coincident with increased FAK phosphorylation, would correlate FAK activity with apoptosis in *in vitro* settings. However, in an animal model of FECD, pharmacological inhibition of FAK led to increased corneal endothelial cell loss. This lack of consistency between *in vitro* and *in vivo* findings prevents us from speculating whether FAK would function in a cell-protective or apoptotic manner in the corneal endothelial cells.

Our goal in this study was to understand how the physical changes to the Descemet’s membrane affect corneal endothelial cell functions. By comparing cell behaviors in soft and stiff substrates, we identified FAK activity to be increased only by the change in stiffness of the substrate. A major limitation of this study is the use of primary bovine corneal endothelial cells in place of human corneal endothelial cells. While the Descemet’s membrane stiffness in normal and disease states is characterized for human samples and mouse models, the normal Descemet’s membrane stiffness for bovine corneas is different.^48^ However, we reasoned that, by growing the bovine cells in plates that vary 4-fold in stiffness, we can compare the relative effect of this parameter on cell health. Furthermore, the use of *ex vivo* and *in vivo* studies in mice bolsters our *in vitro* findings. During disease progression, there are many changes to the Descemet’s membrane, not just in its stiffness. Studying only the effect of stiffness helped us to identify FAK activity as an outcome of this change. Understandably, inhibiting FAK activity alone did not improve all the phenotypes in the *Col8a2^Q455K/Q455K^* knock-in FECD mice, given the multifactorial nature of the disease onset. However, improvement in the levels of antioxidants and a decrease in EndMT provide important information on how ECM stiffness changes may affect corneal endothelial cell functions.

## Data availability

All relevant raw data are available upon request from the corresponding author.

## Conflicts of interest

**SAG:** None, **VSC:** None, **SM:** None, **SH:** None, **RS:** None

## Author contributions

**SAG:** Writing - original draft, investigation, and data analysis. **VSC:** investigation. **SM:** investigation. **SH:** Writing- review & editing. **RS:** conceptualization, resources, Writing- review & editing, supervision, funding acquisition.

## Acknowledgments

We gratefully acknowledge grant support from the National Institutes of Health (R00EY032974 (RS), R01EY034096 and R21EY036189 (SH)), the Cornea Research Foundation (RS), the BrightFocus Foundation G2024006S (SH), Research to Prevent Blindness (David Epstein Career Advancement Award in Glaucoma Research; SH), and unrestricted grants from Research to Prevent Blindness and Lions Region 20-Y1 to SUNY Upstate Medical University Department of Ophthalmology and Visual Sciences (SH).

## Declaration of generative AI and AI-assisted technologies in the manuscript preparation process

During the preparation of this work the author(s) used Grammarly in order to correct grammatical errors. After using this tool/service, the author(s) reviewed and edited the content as needed and take(s) full responsibility for the content of the published article.

---

**Table 1.**
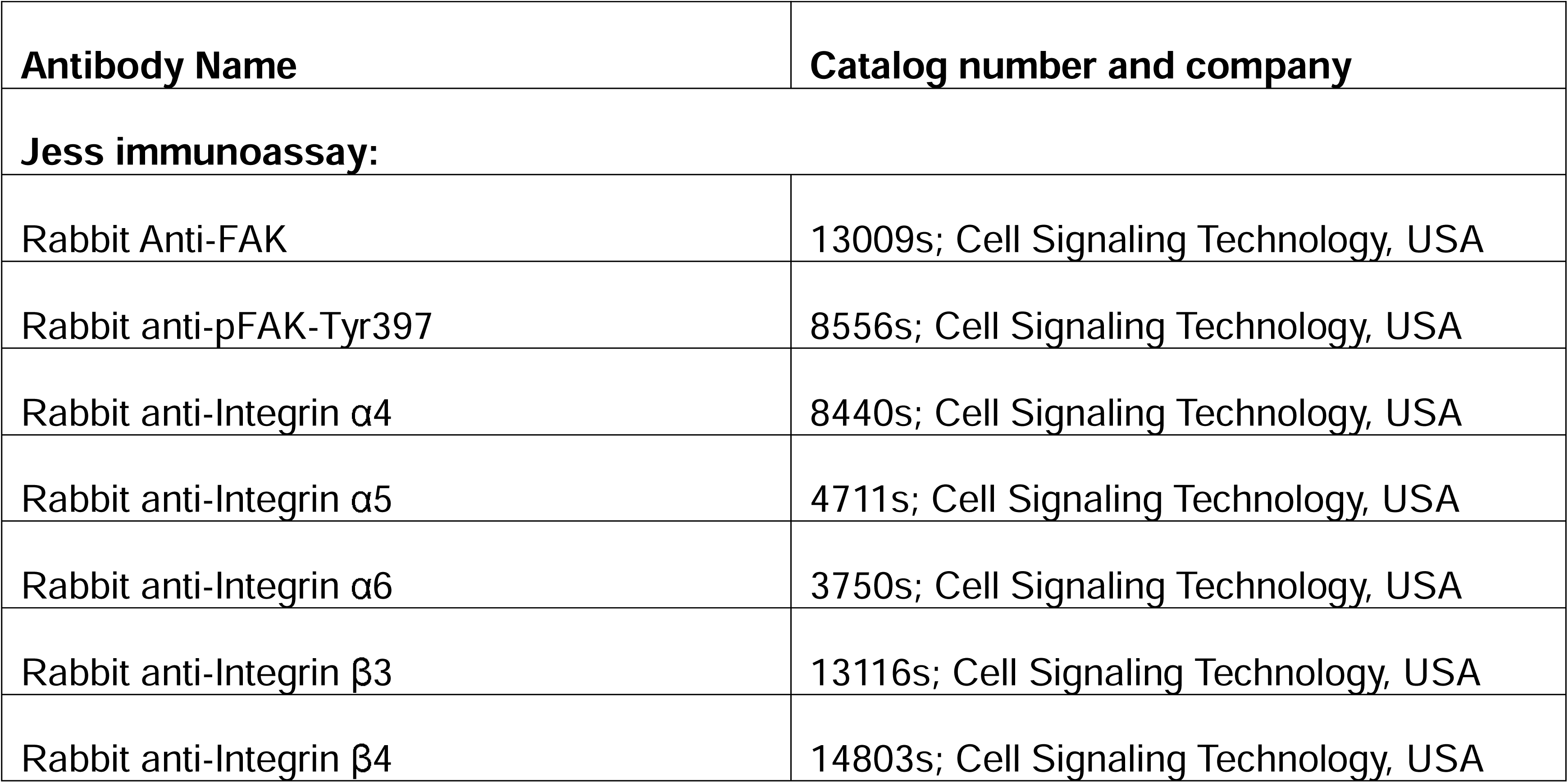
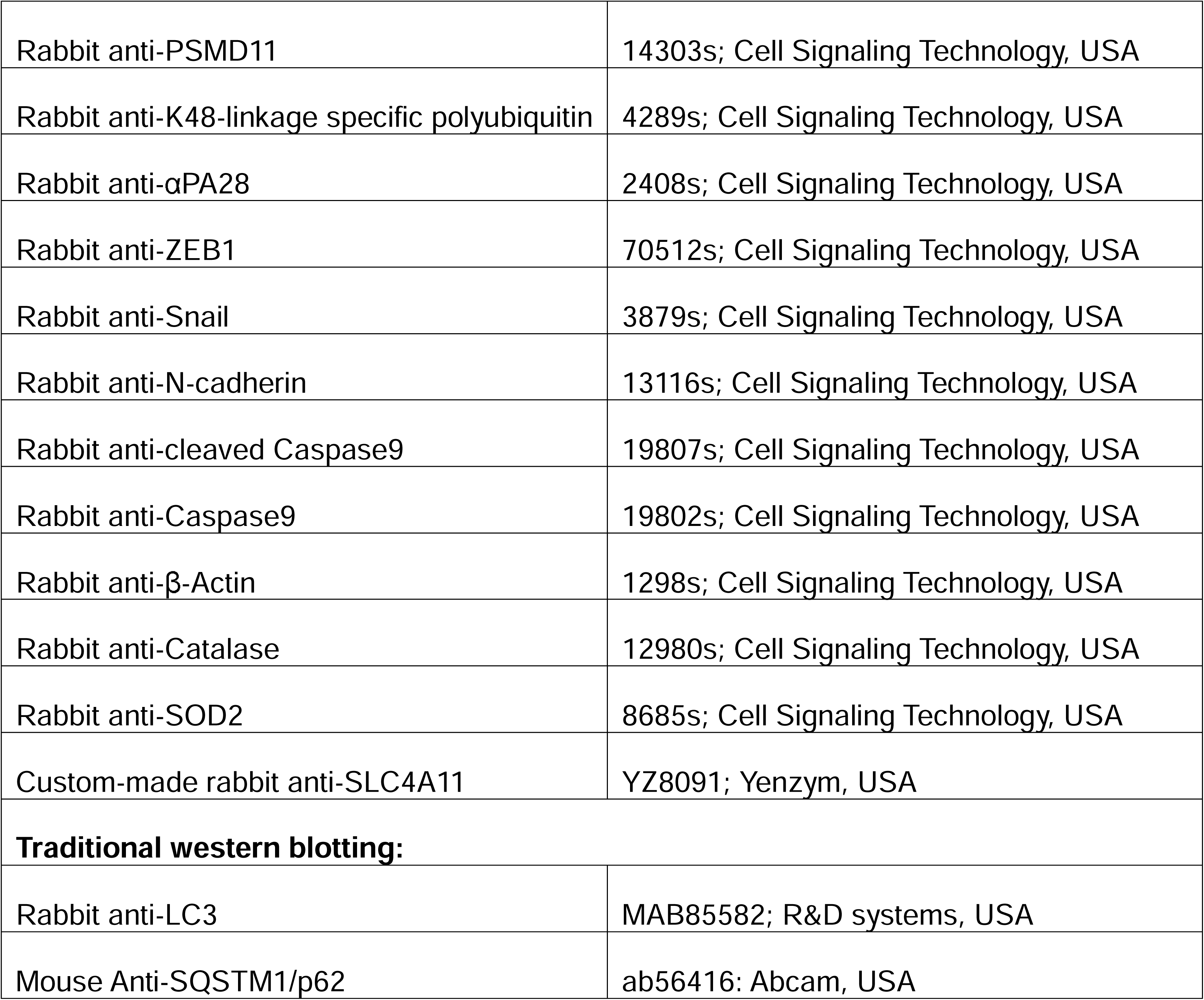
List of antibodies used for Jess immunoassay and traditional western blotting.

| Antibody Name | Catalog number and company |
| --- | --- |
| **Jess immunoassay:** |  |
| Rabbit Anti-FAK | 13009s; Cell Signaling Technology, USA |
| Rabbit anti-pFAK-Tyr397 | 8556s; Cell Signaling Technology, USA |
| Rabbit anti-Integrin α4 | 8440s; Cell Signaling Technology, USA |
| Rabbit anti-Integrin α5 | 4711s; Cell Signaling Technology, USA |
| Rabbit anti-Integrin α6 | 3750s; Cell Signaling Technology, USA |
| Rabbit anti-Integrin β3 | 13116s; Cell Signaling Technology, USA |
| Rabbit anti-Integrin β4 | 14803s; Cell Signaling Technology, USA |
| Rabbit anti-PSMD11 | 14303s; Cell Signaling Technology, USA |
| Rabbit anti-K48-linkage specific polyubiquitin | 4289s; Cell Signaling Technology, USA |
| Rabbit anti-αPA28 | 2408s; Cell Signaling Technology, USA |
| Rabbit anti-ZEB1 | 70512s; Cell Signaling Technology, USA |
| Rabbit anti-Snail | 3879s; Cell Signaling Technology, USA |
| Rabbit anti-N-cadherin | 13116s; Cell Signaling Technology, USA |
| Rabbit anti-cleaved Caspase9 | 19807s; Cell Signaling Technology, USA |
| Rabbit anti-Caspase9 | 19802s; Cell Signaling Technology, USA |
| Rabbit anti-β-Actin | 1298s; Cell Signaling Technology, USA |
| Rabbit anti-Catalase | 12980s; Cell Signaling Technology, USA |
| Rabbit anti-SOD2 | 8685s; Cell Signaling Technology, USA |
| Custom-made rabbit anti-SLC4A11 | YZ8091; Yenzym, USA |
| **Traditional western blotting:** |  |
| Rabbit anti-LC3 | MAB85582; R&D systems, USA |
| Mouse Anti-SQSTM1/p62 | ab56416; Abcam, USA |

## Notes

### Competing Interest Statement

The authors have declared no competing interest.

### Summary of Updates

Corrections of errors in figures and in the manuscript body, and improved resolution of figures.

